# Wide awake at bedtime? The effects of caffeine on sleep and circadian timing in teenagers - a randomized crossover trial

**DOI:** 10.1101/2020.03.06.980300

**Authors:** Carolin F. Reichert, Simon Veitz, Miriam Bühler, Georg Gruber, Sophia S. Rehm, Katharina Rentsch, Corrado Garbazza, Martin Meyer, Helen Slawik, Yu-Shiuan Lin, Janine Weibel

## Abstract

**Background:** Adolescents frequently consume caffeine with unknown consequences on sleep and circadian rhythms. In adults, the evidence indicates that caffeine acutely reduces homeostatic sleep pressure and delays the circadian timing system.

**Objective:** Here, we investigated the acute effects of caffeine intake on the developing sleep-wake regulatory system of teenagers.

**Design:** In a double-blind randomized crossover laboratory study, 18 teenagers (16.1 ± 1 years old, pubertal development scale [PDS]: 2.76 ± 0.35) ingested 80 mg caffeine (vs placebo) four hours prior to bedtime. Until bedtime, participants regularly filled in the Karolinska Sleepiness Scale and gave saliva samples to measure melatonin secretion. During nighttime, we quantified homeostatic sleep need by the electroencephalographically derived amount of slow wave sleep duration (SWS). After sleep, participants rated sleep quality by the Leeds Sleep Evaluation Questionnaire.

**Results:** While participants felt less sleepy after caffeine vs. placebo (*P*=0.038), their ratings of sleep quality were not strongly affected by the treatment. However, objectively, SWS was on average reduced by ∼20 min after caffeine vs. placebo (*P*=0.026). This caffeine-induced reduction was more pronounced in those individuals with more SWS under placebo (regression: *P*=0.042; standardized beta=0.622, *P*=0.011), even if controlling for habitual caffeine intake or pubertal stage (PDS). In melatonin onsets we observed both delays and advances in response to caffeine. This variance could partly be explained by differences in relative dose (i.e., mg caffeine/kg of bodyweight): the higher the relative dose, the more likely were delays (regression: *P*=0.01; standardized beta=0.592, *P*=0.01).

**Conclusions:** In teenagers, evening caffeine intake of already 80 mg (i.e. ∼8fl oz of common energy drinks) is sufficient to promote alertness at the costs of subsequent sleep. These costs might be more pronounced in adolescents with a higher need for SWS. Moreover, caffeine might disturb the circadian timing system, consequently hampering the balanced interplay of sleep-wake regulatory components.

**Sources of Support:** Marie-Heim-Voegtlin Grant of the Swiss National Science Foundation (SNSF) PMPDP1_171364

## Introduction

Around 80% of teenagers consume caffeine (1-3), a psychoactive stimulant which is present in a variety of foods and over-the-counter beverages and medications (4). The amounts of daily intake strongly vary from 40 mg reported by adolescents in the UK up to 350 mg in Austria (5-9). However, due to a lack of empirical studies (10, 11), the consequences of caffeine-intake on the developing neuronal and cardiovascular system are rather unclear. Accordingly, evidence-based limits for safe caffeine consumption in children and adolescents are missing up to date (10, 11) and the American Academy of Pediatrics strongly recommends to eliminate the stimulant from the daily diet in this age group (12). At the same time caffeine-containing beverages, in particular so-called energy-drinks, are aggressively marketed towards teenagers and young adults (13), often promising a boost in physical and mental capacities (13, 14). These claims indeed reach their target population (15), is mirrored in the teenagers’ motivation to consume caffeine-containing drinks in order to enhance performance and reduce sleepiness (1, 16).

As conceptualized in the “perfect storm model” (17, 18), high daytime sleepiness of teenagers can arise from a conflict between social constraints and developmental changes in both sleep-homeostatic and circadian components of sleep-wake regulation (19). While school times often require early rise times, the teenagers’ sleep-wake regulation supports bedtimes late at night, as the increase of homeostatic sleep need slows down (20) and the circadian timing system becomes phase-delayed (21, 22) as compared to childhood. Hence, it is not surprising that in the US around 75% of teenagers do not get the required 9 hours of sleep (23) and around 40% of teenagers suffer from excessive daytime sleepiness (23-25).

As an adenosine receptor antagonist of the central nervous system, caffeine can disturb the homeostatic regulation of sleep need (26). In adults, the acute intake of the stimulant has repeatedly been shown to reduce sleep depth, sleep duration (27) and subjective sleepiness (28). Moreover, recent evidence in adults suggests that caffeine consumption particularly in the evening suppresses melatonin secretion (29, 30) and delays the circadian onset of the biological night (31). While caffeine intake might thus appear at first glance to be well-suited to counteract sleepiness during adolescence, caffeine consuming teenagers report a higher sleepiness compared to non-consumers (32). This could potentially be linked to caffeine-induced disturbances in sleep-wake regulation, as caffeine-consuming teenagers also report a shorter sleep duration (23, 33-36), less restorative sleep (33, 37) and show a reduced depth of sleep (38). However, cross-sectional studies are not suitable to identify cause-effect-relationships and experimental laboratory studies are missing so far. Thus, we examined the consequences of caffeine intake in adolescents using a placebo-controlled double-blind crossover design. We hypothesized that one-time caffeine intake reduces shortens slow wave sleep duration and delays the circadian melatonin onset in teenagers. A secondary objective was to test the assumption that caffeine acutely reduces subjective sleepiness and sleep quality.

## Methods

### Sample Size

The available evidence in adults indicated that caffeine intake in the evening elicits large effects on both slow wave sleep (SWS) duration (assumed d=1.14 on the basis of (39); and d=1.38 on the basis of (40)) and circadian phase (d=0.93, (31)). Thus, accepting α=0.05 and β =.20, a sample size of N=18 could be considered to be sufficient to detect significant caffeine-induced effects of this size when assuming calculation of a t-test for dependent samples (analyses done with G*Power 3.1.9.2., (41)).

### Subjects

Participants were recruited via online platforms at the University of Basel, posts at sports clubs and presentations at schools in Basel, Switzerland, and the surrounding area. Overall, 68 teenagers completed a telephone interview, to check inclusion criteria (14-17 years of age, sex: male, BMI: 16.2-25.4, right handedness, sleep duration during school days: 6-10 h, moderate chronotype [according to the Munich Chronotype Questionnaire (42) [MSF-Sc]: 2-6.99). The latter two criteria were set to avoid abnormal sleep pressure levels and extreme timing of circadian phase during the laboratory part of the study. Individuals were only included if habitual caffeine intake ranged between 80 and 300 mg per week [assessed with a survey tool based on (43) and adapted to the caffeine content according to (44)] in order to avoid naïve and/or highly sensitive individuals. The upper limit of 300 mg/week was set to keep potential withdrawal symptoms at low levels during the week of abstinence preceding the laboratory part. Included participants were asked to fill in questionnaires to quantify pubertal stage (Pubertal Development Scale (PDS) (45)), to exclude chronic diseases (such as diseases of the respiratory or coronary system) and to check on sleep behavior (Pittsburgh Sleep Quality Index (46)). In addition, the physician in charge conducted the neuropsychiatric interview MINI KID (47) in order to exclude participants suffering from psychiatric disorders. In a last step before start of study, participants were invited for an adaption night in the sleep laboratory, to allow habituation to the sleep rooms and electro-encephalographic (EEG) instruments, to explain the study procedures and to conduct a urinary drug screening (Drug-Screen-Multi 6, nal von minden GmbH, Germany) in order to exclude recent drug use.

As illustrated in supplemental Figure 1, of 68 individuals that had been contacted by phone, 15 met one of the exclusion criteria and 28 could not be contacted anymore or lost interest in participation. From the remaining 25 individuals, one had to be excluded due to medical reasons, five dropped out after the habituation night and one dropped out during the first laboratory condition. Thus, the final sample consisted of N=18 participants, completing both conditions. Demographics of this sample are summarized in Table 1.

**Table 1.** Demographic information on participants. Regular sleep duration and regular bedtime were extracted from the Munich Chronotype Questionnaire (42). MCTQ MSF-Sc: Chronotype according to Munich Chronotype Questionnaire; PDS: Pubertal Development Scale (45).

| Variable (unit) | *M* | *SD* |
| --- | --- | --- |
| Age (years) | 16.11 | 1.09 |
| Regular sleep duration at schooldays (h:min) | 7:45 | 36 min |
| Regular bedtime during schooldays (hh:min) | 22:32 | 36 min |
| Bedtime during study (hh:min) | 22:36 | 37 min |
| Body Mass Index (kg/m<sup>2</sup>) | 20.80 | 2.13 |
| Habitual caffeine intake (mg/week) | 248.50 | 122.78 |
| Chronotype (MCTQ MSF-Sc) | 3.80 | 0.78 |
| Pubertal Stage (PDS) | 2.76 | 0.35 |

Please note that the available evidence in adults indicates that caffeine intake in the evening elicits large effects on both slow wave sleep (SWS) duration (assumed d=1.14 on the basis of (39); and d=1.38 on the basis of (40)) and circadian phase-shifts (d=0.93, (31)). Thus, accepting α=0.05 and β =.20, a sample size of N=18 can be considered to be sufficient to detect significant caffeine-induced effects of this size when assuming calculation of a t-test for dependent samples (analyses done with G*Power 3.1.9.2., (41)).

### Study Protocol

The procedures were conducted in accordance with the ethical standards of the responsible regional ethical committee (Ethikkommission Nordwest- und Zentralschweiz) and in accordance with the Helsinki Declaration of 1975 as revised in 1983. Every participant and one legal representative of each participant signed informed consents.

Every participant took part in two conditions (caffeine and placebo), which were separated on average by 8.5 days (range: 6-35 days, median: 7 days). The order of conditions (caffeine vs placebo) was randomized (55% of the sample had placebo first, see supplemental Figure 1). Salivary caffeine and paraxanthine values at outset did not significantly differ between these two groups (*P*>0.5) indicating similar starting points and a sufficient washout period between conditions. As illustrated in supplemental Figure 2, both caffeine and placebo conditions started with an ambulatory period of six days, to control for sleep debt and allow circadian entrainment as well as washout from prior habitual caffeine intake. To do so, participants were asked to adhere to a fixed sleep-wake cycle (8 h sleep window, no naps allowed, allowed deviation from bed- and waketime ± 60 min) and abstain from all caffeine-containing foods and beverages. Sleep and wake times for the ambulatory weeks were set individually according to the individual preferences and social constraints (i.e., school times) and compliance was checked by means of wrist actigraphy (Actiwatches®, Cambridge Neurotechnology Ltd., Cambridge, UK). Salivary caffeine levels and levels of the main caffeine metabolite paraxanthine (depicted in Figure 1) were checked at outset of the laboratory parts.

**Figure 1.**
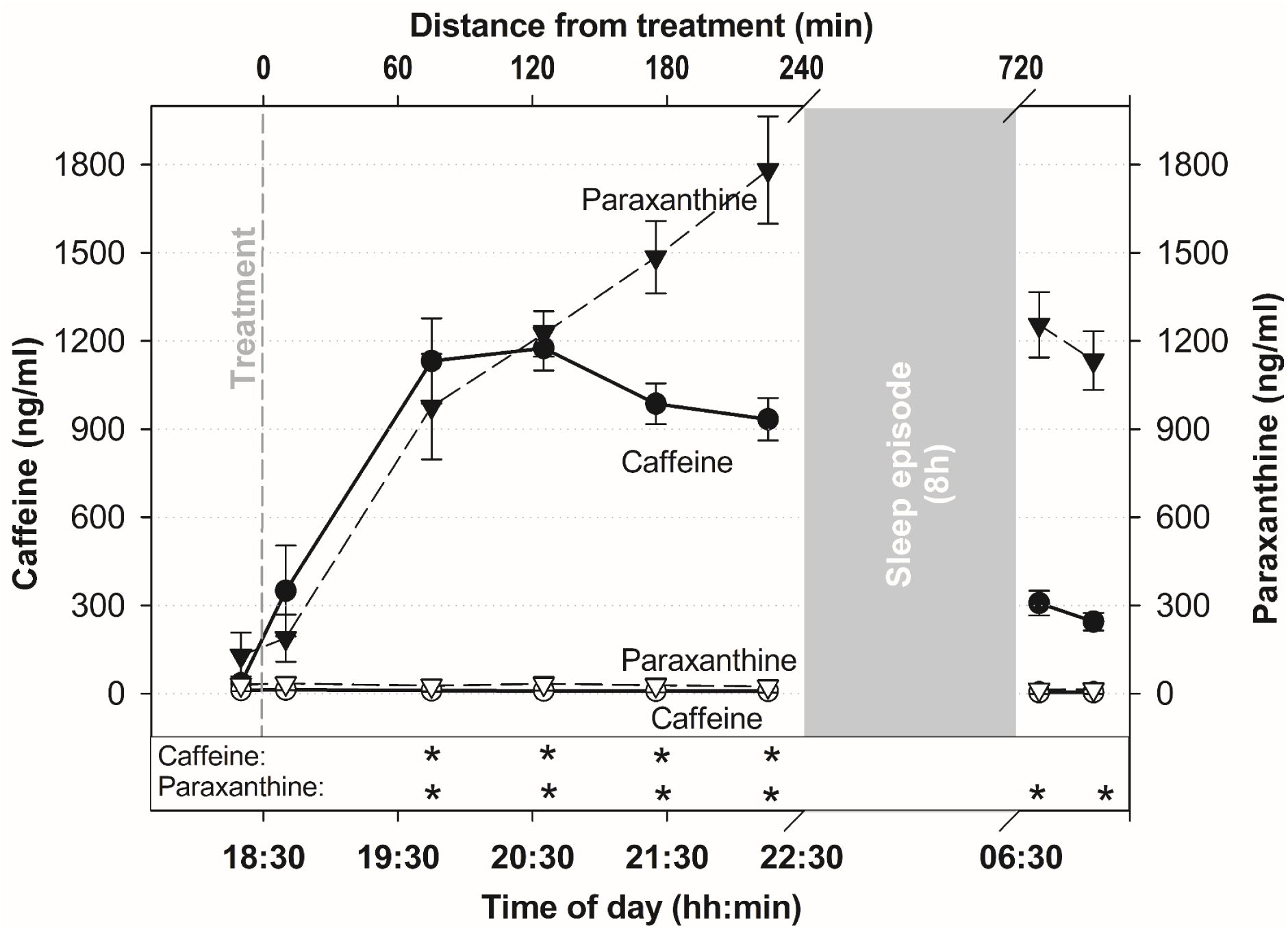
Course of salivary caffeine and paraxanthine levels per condition. Black symbols indicate the course of caffeine (circles) and paraxanthine (triangles) during the caffeine condition, while white symbols indicate the levels (caffeine: circles, paraxanthine: triangles) assessed during the placebo condition. Time-of-day values on the x-axis represent group means, as time of treatment was adjusted to the individual bedtime of each participant. Stars indicate significant differences between conditions per time of measurement for caffeine and paraxanthine, respectively (*P* <0.05, adjusted for multiple comparisons).

Both laboratory parts started in the evening 5 h before habitual sleep time (*M*: 10:36 p.m., *SD*: 37 min) and did only vary with regard to the treatment. The caffeine or placebo capsule was given 4 h before habitual bedtimes and contained either 80 mg of caffeine and mannitol or mannitol only (Hänseler AG, Herisau, Switzerland). The dose of 80 mg roughly equals not only the daily average intake of adolescents in the US (3, 5) but also the content per serving in a range of common energy drinks (e.g., Bomb Energy Drink® (8.45 fl oz), Burn® (8.3 fl oz), Hell Energy® (8.4 fl oz), Monster® Regular (Green) (8 fl oz), NOS® (8 fl oz), Pussy® (8.45 fl oz), RedBull® (8.45 fl oz), Rockstar® Energy Drink (8 fl oz), etc.). We set the timing of administration to the evening hours to allow maximal effects of caffeine on both sleep (39, 48, 49) and circadian rhythms (31, 50). Please note that teenagers represent the age group in which evening caffeine consumption is most common (5).

### Subjective Sleepiness and Sleep Quality

During scheduled wakefulness, subjective sleepiness was assessed every 45 min with the Karolinska Sleepiness Scale (KSS, (51)), a scale frequently used in teenagers to measure the sleepiness response to sleep-wake manipulations such as total (52) or partial sleep deprivation (53-55). Volunteers rated their sleepiness within the past 10 min using a 9 point verbally anchored scale from 1 (extremely alert) to 9 (extremely sleepy, fighting sleep). Due to technical problems, we lost one data set in the caffeine condition.

To assess subjective sleep quality, we used the Leeds Sleep Evaluation Questionnaire (LSEQ, (56)) administered in the morning right at wakeup. The evaluation of the questionnaire results in four dimensions *Getting to Sleep, Quality of Sleep, Awake Following Sleep* and *Behavior Following Wakening* (57). Due to technical problems, we lost the data of one participant in the caffeine condition.

Expectations might particularly influence subjective ratings of sleepiness and sleep. However, those participants who correctly identified the caffeine condition at the end of the study, did not significantly differ on average in ratings of sleepiness or sleep quality (*P*>0.14) compared to the other participants.

### Salivary Caffeine and Melatonin

We took saliva samples every 45 min during scheduled wakefulness under dim-light conditions. Caffeine levels as well as levels of the main caffeine metabolite paraxanthine were determined with liquid chromatography coupled to tandem mass spectrometry. The area under the curve (AUC) of caffeine levels in the evening during the caffeine condition was calculated according to a trapezoid formula described in (58), taking baseline levels before caffeine treatment into account (i.e., AUC_I_).

Melatonin levels were measured using a direct double-antibody radio immunoassay (59). Dim-light melatonin onset (DLMO) was quantified using the hockey-stick method (60) with an ascending level of 2.5 pg/ml. We calculated the AUC of melatonin levels in the evening with respect to the ground (i.e., AUC_G_) using the trapezoid formula described in (58).

### Sleep EEG

For recordings of sleep electroencephalogram (EEG), electrooculogram (EOG) and electromyogram (EMG), we used Live-Amp devices (Brain Products GmbH, Gilching, Germany) in combination with electrode caps (32Ch LiveAmp Cap with Multitrodes, Easycap GmbH, Herrsching, Germany). Each of the two nights of every participant was recorded with the same device. We recorded signals with a sampling rate of 250 Hz, applying an online notch filter at 50 Hz, at derivations from the frontal, central, parietal and occipital regions (F3, FZ, F4, C3, CZ, C4, P3, PZ, P4, O1, OZ and O2) referenced against FCz. Offline the signals were re-referenced against the contra-lateral mastoid (A1 or A2) according to the guidelines of the American Academy of Sleep Medicine (AASM, https://aasm.org/clinical-resources/scoring-manual/) and downsampled to 128 Hz for automatic sleep staging by a validated (61) algorithm (Somnolyzer 24 × 7, The Siesta Group, Vienna, Austria). Scorings were visually controlled epoch-by-epoch according to the AASM criteria (62) by an expert scorer (and all automatic scorings were considered as correct). This approach combines the reliability of standardized automatic scouring with the validity of human expert scoring and has successfully applied in earlier studies on sleep of adolescents (e.g. (63, 64)). Due to technical problems, the recordings of two nights (1 x placebo condition, 1 x caffeine condition) were incomplete. Thus, these data were not included into the analyses but replaced by the means of each parameter across the group and the two conditions.

### Statistical Analyses

To analyze KSS, salivary caffeine and salivary paraxanthine values, we applied mixed models using the SAS 9.4 software (SAS Institute, Cary, USA). To assess both differences between conditions (caffeine vs placebo) and condition-specific differences in time courses (interaction between factors condition and time) within one model we used the procedure PROCMIXED. We included subject as a random factor and assumed the covariance structure AR(1) (i.e., autoregressive 1) for the factor time. P-values are based on corrected degrees of freedom according to Kenward and Roger (65). Post-hoc comparisons were calculated using the LSMEANS statement and were adjusted for multiple comparisons according to the Tukey-Kramer method.

Using the software package SPSS (Version 25, IBM Corp., Armonk, USA), we compared subjective sleep quality, proportion of SWS (percentage of total sleep time) and DLMO, calculating t-tests for dependent measures. If not otherwise indicated, we report *P*-Values based on one-sided testing as we tested directed hypotheses.

To investigate predictors for caffeine-induced responses in sleep need and circadian timing, we calculated two regression models separately to explain a) the caffeine-induced difference in SWS proportion (i.e., SWS proportion placebo - SWS proportion caffeine) and b) the caffeine-induced difference in DLMO (i.e., DLMO in caffeine - DLMO in placebo condition). Collinearity between predictors was considered as substantial if predictors were correlated at *p*<0.05, one-sided. Regarding a), the following possible predictors were included: relative caffeine dose ((66, 67) quantified by mg/kg body weight), actual state of caffeine metabolism (assessed by the ratio paraxanthine/caffeine in saliva (68) taken 15 min before start of sleep episode in the caffeine condition) and an approximation of individual SWS need (i.e., SWS proportion during the placebo night). Due to collinearity with the latter predictor, we did not include self-reported habitual intake levels (assessed by (43)), age or pubertal stage (assessed by PDS (45)) in the final regression model. Regarding b), we aimed at exploring reasons for our results diverging from our hypothesis and earlier studies in adults (29-31). Thus, the following predictors were included into a stepwise backward regression: age, relative given dose (quantified by mg/kg body weight) and habitual intake levels (assessed by (43)). Pubertal stage (assessed by PDS (45)) and distance of treatment to DLMO were not included due to collinearity.

## Results

### Salivary Caffeine

Figure 1 shows the course of caffeine and paraxanthine levels over time in both conditions. Before treatment, caffeine levels in the caffeine condition did not differ from those in the placebo condition, but were significantly increased from 75 min after intake until bedtime compared to the placebo condition (*F*_[7,135]_=22.89, *P*<0.001). Similarly, paraxanthine levels (i.e., levels of the main metabolite of caffeine) did not differ between conditions before treatment but were increased in the caffeine condition starting from 75 min after intake until the end of the study compared to the placebo condition (i.e., 12,7 h after intake (*F*_[7,92.5]_=17.17, *P*<0.001)).

### Subjective Sleepiness and Sleep Quality

As depicted in Figure 2A, on average participants indicated lower sleepiness values in the caffeine compared to the placebo condition (*F*_[1,40.9]_=4.58, *P*=0.038; *M*_caff_±*SD*=4.38±1.92, *M*_plac_±*SD*=4.78±2.01). However, the increase of sleepiness towards bedtime (*F*_[7,108]_=9.59, *P<*0.001) did not significantly differ between conditions (*F*_[7,116]_=1.85, *P*=0.084; Figure 2B).

**Figure 2.**
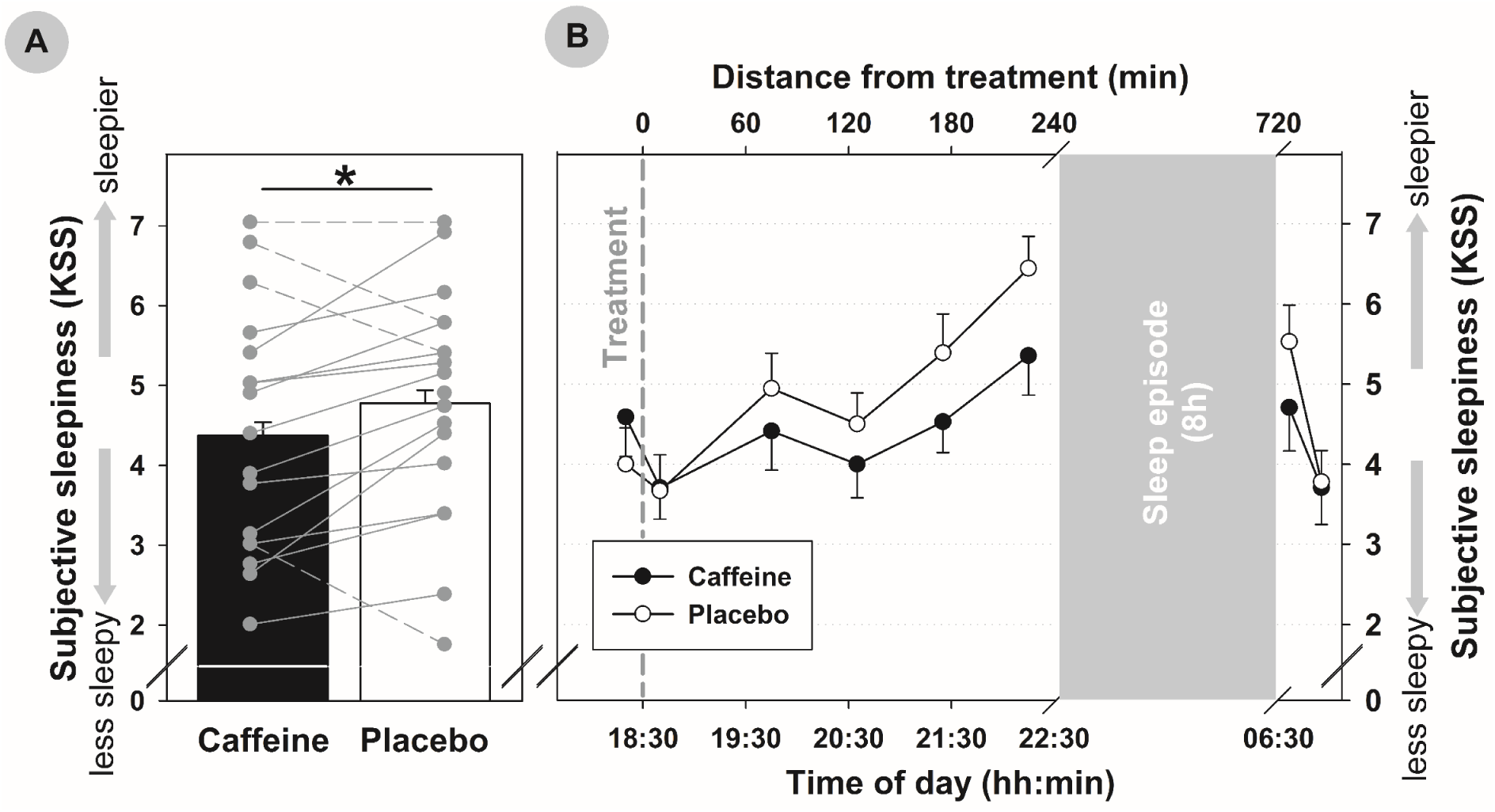
Differences in subjective sleepiness between conditions. A) Subjective sleepiness (as assessed by the Karolinska Sleepiness Scale, KSS (51)) was on average lower (*: *P*<0.05) in the caffeine (black bar, mean + standard error) compared to the placebo condition (white bar, mean + standard error). This caffeine-induced reduction of sleepiness was present in ∼75% of individuals (solid lines). Other cases are indicated by dashed lines. B) The typical increase in subjective sleepiness towards bedtime is shown as group mean with standard error per time of assessment. The increase was not significantly modulated by condition (interaction time x condition: *P*=0.084). As time of treatment was adjusted to the individual bedtime of each participant, time-of-day values on the x-axis represent group means.

The analyses of subjective sleep quality revealed a significant difference for the scale *Getting to Sleep* only (*t*_(16)_=-2.16, *P*=0.023; *M*_caff_±*SD*=40.95±14.49, *M*_plac_±*SD*=43.38±16.75), indicating that participants noticed more problems with falling asleep in the caffeine condition compared to placebo. The other scales (i.e., *Quality of Sleep, Awake Following Sleep* and *Behavior Following Wakening*) did not significantly differ between conditions.

### Sleep Patterns

The proportion of all sleep stages and sleep latencies during the placebo and caffeine condition are summarized in supplemental Table 1. As hypothesized, SWS proportion was reduced after caffeine compared to placebo (*t*_(17)_=-2.093, *P*=0.026; *M*_caff_±*SD*=36.91±8.69, *M*_plac_±*SD*=40.51±11.41, Figure 3A) by 3.6% on average, as mirrored in ∼21 min less time spent in SWS in the caffeine condition compared to placebo. To explain the variance of this reduction (see Figure 3A for individual values), we conducted a regression analysis which revealed that caffeine-induced reductions in SWS proportion (i.e., the individual difference in SWS proportion between placebo and caffeine) were significantly predicted (corrected *R*^2^=0.321, *P*=0.042) by the individual’s SWS proportion during placebo (*β*=0.398, standardized *β*=0.622, *P*=0.011, see Figure 3B for illustration), but not by relative dose (quantified by mg/kg body weight, *P*=0.901) or state of caffeine metabolism right before sleep (ratio of paraxanthine/caffeine, *P*=0.572). Due to collinearity, self-reported habitual intake, pubertal stage or age were not included into the regression model (correlation SWS proportion during placebo night and habitual intake: *r*=-559, *P*=0.016, two-sided; pubertal stage: *r*=-0.494, *P*=0.037; age: *r*=-0.675, *P*=0.002). Importantly, partial correlations indicate that the influence of these collinear factors did not abolish the association between placebo SWS proportion and caffeine induced-differences in SWS (partial correlation controlling for habitual intake *r*=0.538, *P*=0.026, two-sided; for pubertal stage *r*=0.524, *P*=0.031, two-sided; for age *r*=0.552, *P*=0.022, two-sided). Taken together, the individual’s caffeine-induced change of SWS proportion was significantly associated with SWS proportion during placebo, independent of the relative caffeine dose, individual state of metabolism, pubertal stage, age and habitual caffeine intake. The higher SWS proportion was during placebo, the stronger were caffeine-induced changes in this parameter.

**Figure 3.**
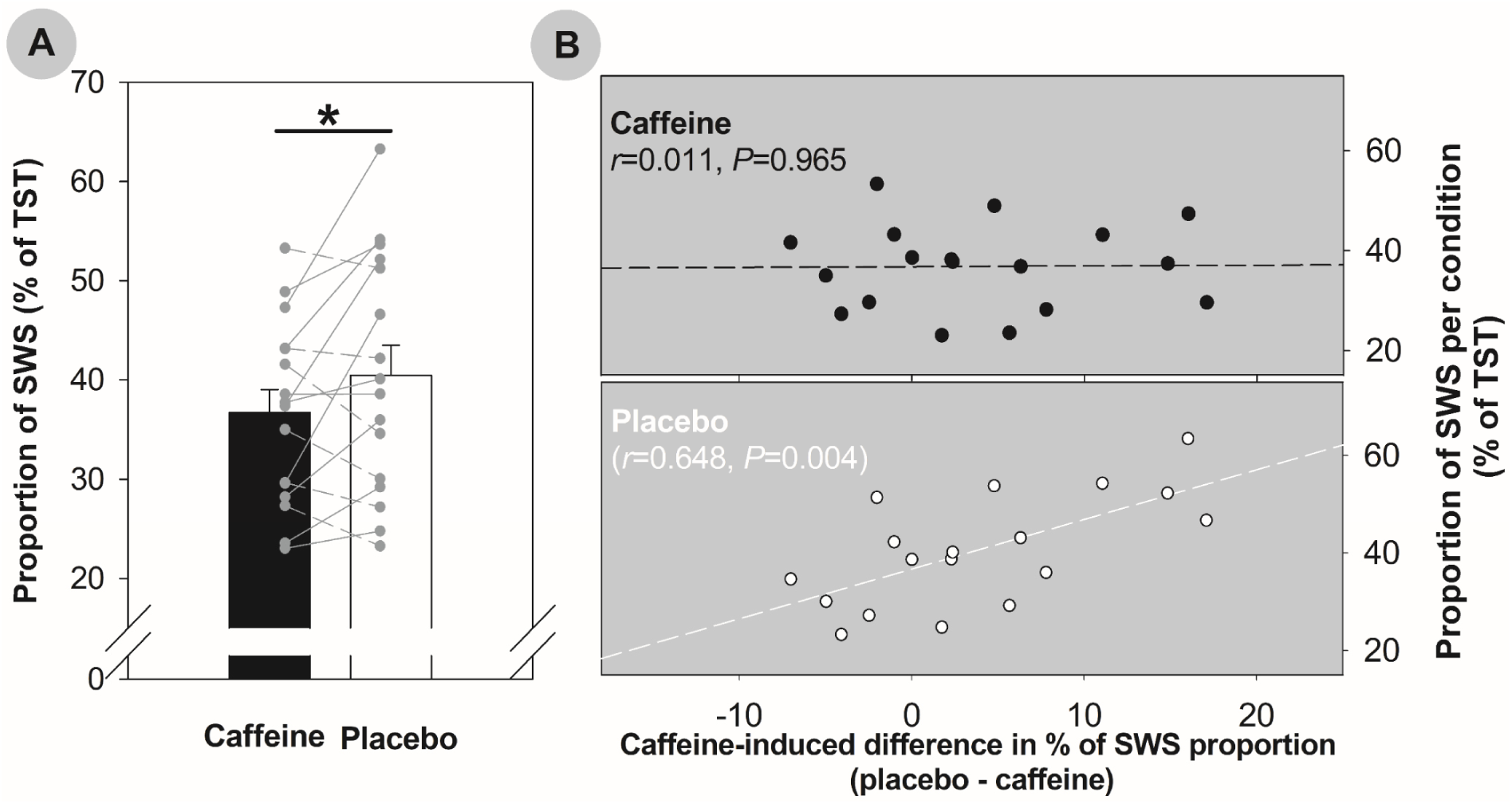
Proportion of slow wave sleep (SWS) per condition and its relation to caffeine-induced changes in SWS. A) The proportion of SWS (as percentage of total sleep time, TST) was significantly reduced in the caffeine condition compared to placebo (*: *P*=0.026, t-test). The proportion of SWS per condition is depicted both as group mean plus standard error and per individual. Solid lines indicate individuals with reductions of SWS proportion, dashed lines indicate individuals with increases in caffeine compared to placebo. B) Association of SWS proportion per condition with difference in SWS proportion between conditions. While there was no association in the caffeine condition (upper panel), the SWS proportion in the placebo condition was positively associated with caffeine-induced changes of this variable (lower panel). That is, the higher the SWS proportion was during the placebo condition, the higher was the caffeine-induced reduction in the caffeine condition. Lines represent correlations; *P*-values are based on two-tailed tests.

### Melatonin

The timing of dim-light melatonin onset (DLMO) was detectable in all subjects in both conditions and showed the typical (21, 69) maturational delay (positive correlation of DLMO and PDS, *r*=0.701, *P*=0.001) in the placebo condition. However, in contrast to our hypothesis, the analysis did not reveal a significant delay in DLMO in the caffeine compared to the placebo condition (*t*_(17)_=-0.189, *P*=0.43; *M*_caff_±*SD*=20:35±46min, *M*_plac_±*SD*=20:37±43min). Similarly, the AUC did not differ between conditions (*t*_(16)_=0.827, *P*=0.420; *M*_caff_±*SD*=894.94±693.85 pg*min/ml, *M*_plac_±*SD*=825.46±723.37 pg*min/ml). As illustrated in Figure 4, there was a high inter-individual variance in the response to the treatment, as mirrored in a similar frequency of observed delays (in 44% of the sample) and advances (in 56% of the sample) of melatonin onsets after caffeine compared to placebo. Thus, we explored potential variables explaining this variance as well as the contradiction to the previously reported results in adults (29, 31). A backward linear regression model (corrected *R*^2^=0.310, *P*=0.010) indicated that of those factors, differing with regard to earlier studies, only the relative dose predicted caffeine-induced shifts in DLMO (*β*=2.226, standardized *β*=0.592 *T*=2.939, *P*=0.010, see Figure 4 for illustration), while age and habitual intake levels did not significantly contribute to caffeine-induced shifts of DLMO (*P*>0.4 and *P*>0.9, respectively).

**Figure 4.**
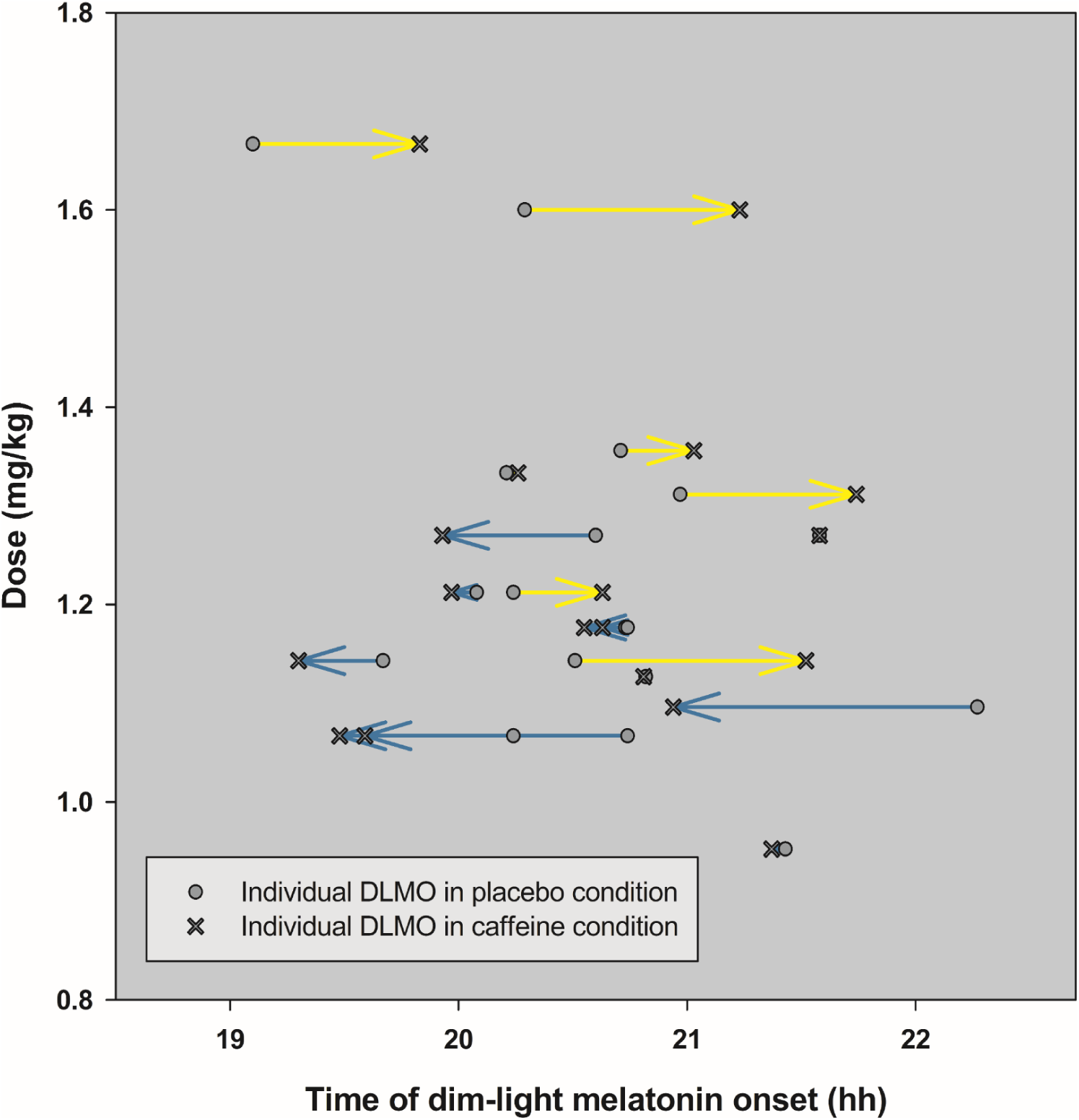
Timing of melatonin onset per participant and condition. Each individual’s dim-light melatonin onset (DLMO) is depicted for the placebo (as circle) and caffeine condition (as cross), according to the relative given caffeine dose on the y-axis. Arrows indicate the shifting direction of DLMO from the placebo towards the caffeine condition within the same individual (yellow arrows represent a delay, blue arrows an advance in DLMO after caffeine intake compared to placebo).

## Discussion

Caffeine is a strong sleep-wake-regulator (27, 70) and part of the daily diet of most teenagers (1-3). The present study is the first to investigate the effects of caffeine on sleep and circadian timing during adolescence using a placebo-controlled randomized crossover design. Our data show that 80 mg of the stimulant in the evening are sufficient to elicit the typical wake-promoting effects at the costs of subsequent sleep. More specifically, caffeine intake reduced feelings of sleepiness, and thus meets one of the main motivations to consume the substance (1). However, it also suppressed SWS during nighttime, i.e. a sleep feature characterized by slow cortical oscillations that are crucial for neuronal recovery (71) and brain maturation (72). Critically, the degree of this caffeine-induced SWS reduction was more pronounced in individuals with relatively high SWS proportion during the placebo night. Thus, caffeine may be particularly sleep-disruptive in those teenagers with a higher sleep need. Finally, in the present sample caffeine intake did delay melatonin secretion as has been observed earlier in adults (29-31), potentially due to the comparably low dose of caffeine used in the present study. Together, the results empirically show the acute stimulating potential of caffeine and thereby underline the need to investigate long-term consequences of the drug on the developing sleep-wake regulatory system.

Already 80 mg of caffeine in the evening, i.e. the dose in 8 fl oz of common energy drinks, were sufficient to effectively promote alertness in teenagers both at a subjective and objective level. While caffeine thus provides an immediate feeling of alertness, the reduced SWS indicates a deficit to recover from the burden of wakefulness during nighttime. This deficit could explain previous observations (25, 73, 74) that daily caffeine intake is generally associated with higher daytime sleepiness as assessed for different daily life situations without specifying if caffeine has been consumed under these circumstances (25, 73, 74). In comparison, our participants rated sleepiness in regard to the past minutes and under the acute influence of the drug. Hence, the diverging results can be reconciled within the assumption that caffeine acutely counteracts sleepiness but that this wake-promoting property might vanish if consumed repeatedly or even express as a withdrawal-related higher subjective sleepiness occurring within a certain time after last intake, as has been shown in adults (75, 76). Given that one of the main reasons for caffeine intake amongst teenagers is to combat sleepiness and enhance alertness (1), caffeine-induced differences in these variables between acute on-time and habitual daily caffeine intake represents an important target in the future regarding both research and education.

In line with a decrease in subjective sleepiness, we observed a reduced amount of SWS after caffeine intake, a sleep stage characterized by cortical slow oscillations and a high slow-wave-activity (SWA) (62). SWA in turn mirrors brain maturation: The decrease in SWA across childhood and adolescence correlates with a decrease in grey matter volume (77, 78), cortical thickness (78) and changes in the grey/white matter ratio (77). Thus, it is possible that reducing SWS by caffeine interferes with the expression of SWA and potentially as well with the related processes of brain maturation. Accordingly, chronic caffeine treatment in pubertal rats delayed the developmental course of SWA and structural markers of brain maturation (79). While consequences of long-term caffeine intake on human brain maturation during adolescence remain to be established, data in human adults indicate that chronic caffeine consumption may indeed lead to a reduced grey matter volume in several cortical regions (80). Taken together, a caffeine dose of 80 mg (i.e. the average daily intake of teenagers in the US (3)) can already impact on sleep structure and disturb SWS. Thus, as long as experimental long-term or longitudinal studies in human adolescents are missing, teenagers may be advised to limit caffeine intake to a minimum.

On the other hand it is important to note that the acute caffeine-induced decrease in SWS by around 20 minutes observed in the present study seems to be rather low as compared to reductions reported in clinical samples (e.g.,(81, 82)) independent of caffeine. Still, together with a caffeine-induced increase in the amount of stage N1 (see supplemental Table 1) during the nighttime sleep episode, our data indicate that the given dose induced rather light and superficial sleep. These differences in sleep structure might be one reason for the worse subjective sleep quality of caffeine-consuming adolescents in the present as well as in other studies (27, 33, 37). Such subjective impressions of bad sleep could not only drive the motivation to in turn combat fatigue by caffeine (1) but also lead to chronic intake and potential long-term adaptations in sleep (38, 79).

Importantly, caffeine intake might particularly affect the homeostasis of sleep in those individuals with a (trait-like) high need for sleep and a higher sleep pressure. For instance, adults show stronger caffeine-induced changes in waking EEG and vigilance the more sensitive they are for high sleep pressure (83). Complementary, our data revealed that the acute amount of caffeine-induced SWS reduction is dependent on the individual amount of SWS during a drug free (i.e., a placebo) night. Although this relation could not be explained by variance from other factors such as pubertal stage, age, or habitual caffeine intake, one might criticize the predictive potential of the SWS amount under placebo as a trait-like indicator for sleep need. For instance, inter-individual variations of SWS duration are only moderately determined by genetic factors (84) and only to some extent stable between conditions. In fact, the amount of SWS is strongly dependent on recent sleep-wake history, such as the duration of prior wakefulness (20) and accumulated sleep debt (85), mirroring the actual need for SWS. However, in the present study, we aimed at controlling the latter by implementing a fixed sleep-wake schedule for six days before start of each laboratory condition. Although limited by a rather small sample size, the present data strongly suggest to further examine whether caffeine-induced sleep-disruptions are particularly strong in populations characterized by a higher need for SWS (e.g., in children), thereby identifying individuals particularly vulnerable for the stimulating potential of the drug.

This need for SWS does not only decrease during the course of adolescence (86-92), but might also vary with habitual caffeine intake. While our data suggest a negative relation between the amount of SWS and the amount of habitual caffeine intake, earlier evidence showed no differences in the amount SWS between habitually caffeine-consuming teenagers compared to non-consumers of the same age (38). Importantly, in the latter study, sleep was recorded under conditions of regular intake while our participants were asked to abstain from caffeine for one week. Keeping in mind that this week of abstinence might not have been sufficient to completely wash out consequences of prior intake, it remains an open question whether impaired SWS is not only a consequence but might also be a risk factor for caffeine intake during adolescence.

Beside the individual need for SWS, we also tested whether the individual caffeine-induced reduction of SWS is related to the individual state of caffeine metabolism. We focused on the ratio paraxanthine/caffeine, a reliable indicator for CYP1A2 activity (68), which in adults metabolizes a large proportion of the caffeine ingested (93) and is most likely not fully developed before the end of puberty (94). However, in our data, metabolic state was not related to the caffeine-induced SWS reduction during nighttime. Together with the available data in human adults (39, 95), the data at present thus do not support a strong predictive power of the absolute metabolic ratio paraxanthine/caffeine on caffeine-induced changes in sleep, but rather suggest a differential adenosinergic signal transduction to be involved in the individual caffeine-induced reduction of SWS (e.g., differential sensitivity of receptors (96)). Such differences might also contribute to inter-individual differences in SWS under placebo conditions (97, 98), which were closely associated with the caffeine-induced SWS response to caffeine.

In contrast to earlier reports, we did not observe a clear shift in the circadian onset of melatonin secretion, considered as the “key to the nocturnal gate of sleep” (98). At first glance, the data do not suggest that evening caffeine intake of 80 mg exacerbates the circadian delay of teenagers (21) in any case. However, apart from the group mean, the results also indicate that higher doses have indeed the potential to do so. In fact, studies in adults indicating a caffeine-induced phase delay (31) or evening melatonin suppression (29, 30) utilized doses which were more than two-fold higher as in the present design (i.e., 2.9 mg/kg in (31) and ∼2.65 mg/kg in (29) vs. ∼1.2 mg/kg in the present study). Moreover, a lower relative dose was already considered as one reason for a weakening in the suppressive effect of evening caffeine on melatonin secretion in adult women (30). However, caffeine-induced phase-advances at low-dose as observed in the present study, have to our knowledge not yet been reported and remain to be replicated. Moreover, it remains to be elucidated which component within the human circadian timing system is affected by caffeine and majorly regulates caffeine-induced shifts in melatonin secretion. A blockade of adenosine receptors might change light responses at the level of the retina (99) or the central circadian master-clock (50, 100) or may even affect melatonin release at the pineal gland (101).

Despite of our rigid study design, this study bears some limitations. First, we included male teenagers only in order to avoid increased variance of sleep and circadian rhythms induced by the menstrual cycle (102) and by the use of oral contraceptives on caffeine metabolism (103, 104). Second, we have no data in an adult control group to specify whether the reported effects of caffeine are specific for maturation during adolescence. In order to approach a potential maturational influence we used pubertal development scores (PDS) and age as continuous variables and explored whether they systematically vary with caffeine-induced responses in the dependent variables. Third, with the rather small samples size we took a risk for false positives and to overlook small effects, such that the results require replication in larger samples. Please note however, that we were interested in large effects and sample size calculations were based on the according effect sizes reported in earlier studies. Fourth, we administered caffeine in the evening hours such that the results might not be generalizable to other timings of intake. Teenagers in the US report to consume a substantial proportion of caffeine specifically in the evening hours (5), and the present data suggest that this intake close to bedtime decreases EEG-derived sleep quality. However, the effects might not necessarily be observed, when caffeine is consumed during the course of the day. In adults for instance, caffeine effects on SWS duration (39) were not detectable anymore when intake was scheduled to the morning hours (48).

In conclusion, the results are the first to empirically show the janus-faced wake-promoting effects of evening caffeine intake in teenagers particularly in the homeostatic process of sleep-wake regulation. The effects were characterized by the typical reduction in subjective sleepiness at the costs of nighttime sleep quality after a relatively low dose of the stimulant. As these costs were higher in teenagers expressing more deep sleep under baseline conditions, our data raise the question whether caffeine could be particularly harmful in individuals with a high homeostatic sleep need, such as children. As long as further empirical studies on consequences and underlying adenosinergic mechanisms are missing, caffeine should thus be consumed with caution.

## Acknowledgements

We thank Claudia Renz for the analyses of melatonin and Michael Strumberger for his contributions to the EEG measurements. We also thank the teenagers for participation and compliance as well as their parents for support.

## Authors contributions

CFR designed research (project conception, development of overall research plan, and study oversight); CFR, SV, MB, CG, HS, MM conducted research (hands-on conduct of the experiments and data collection); CFR, SSR, KR, GG analyzed data or performed statistical analysis; CFR, Y-SL, JW wrote paper (only authors who made a major contribution); CFR, JW had primary responsibility for final content.

## Supplemental Material

**Supplemental Figure 1.**
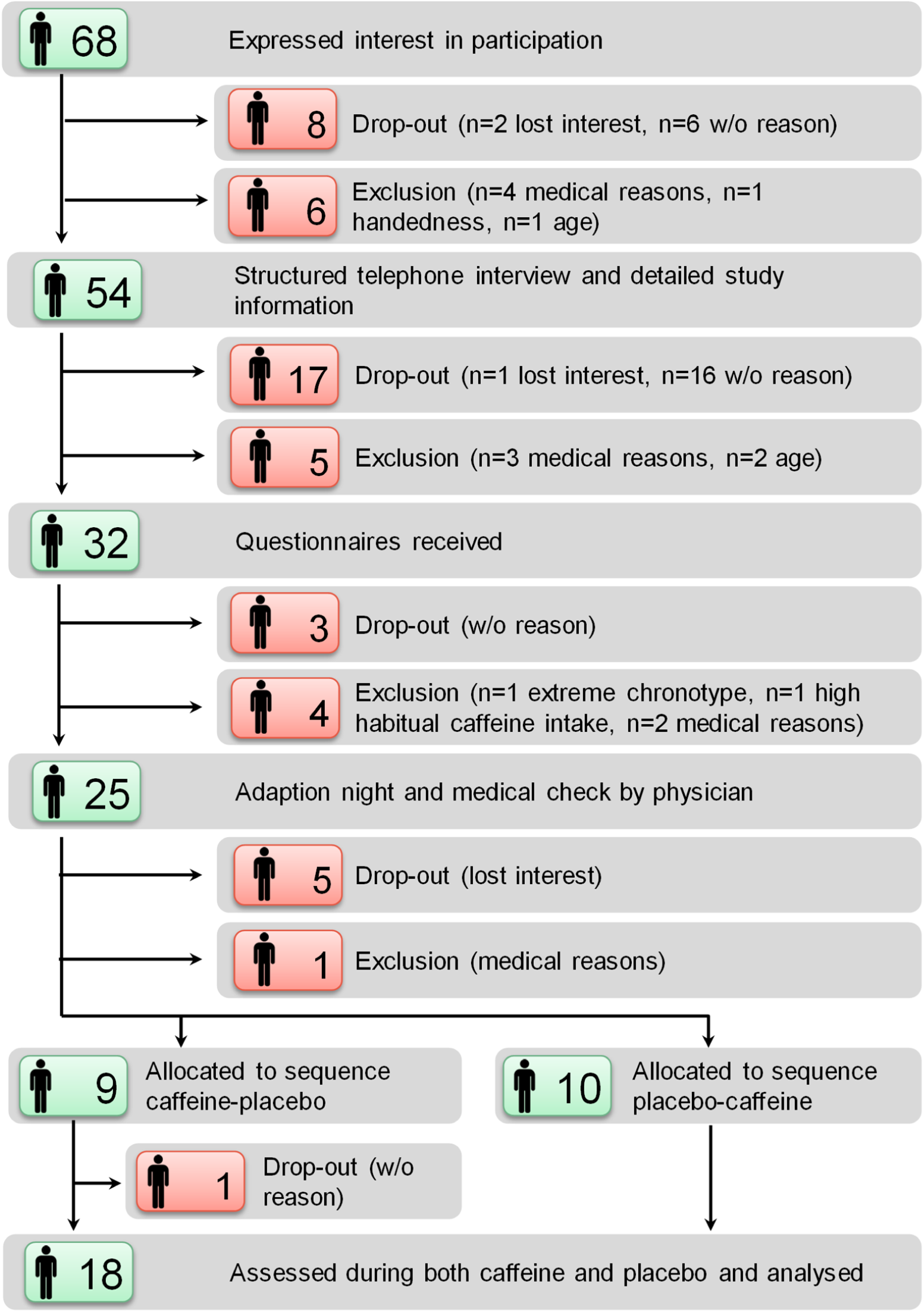
Illustration of recruitment flow.

**Supplemental Figure 2.**
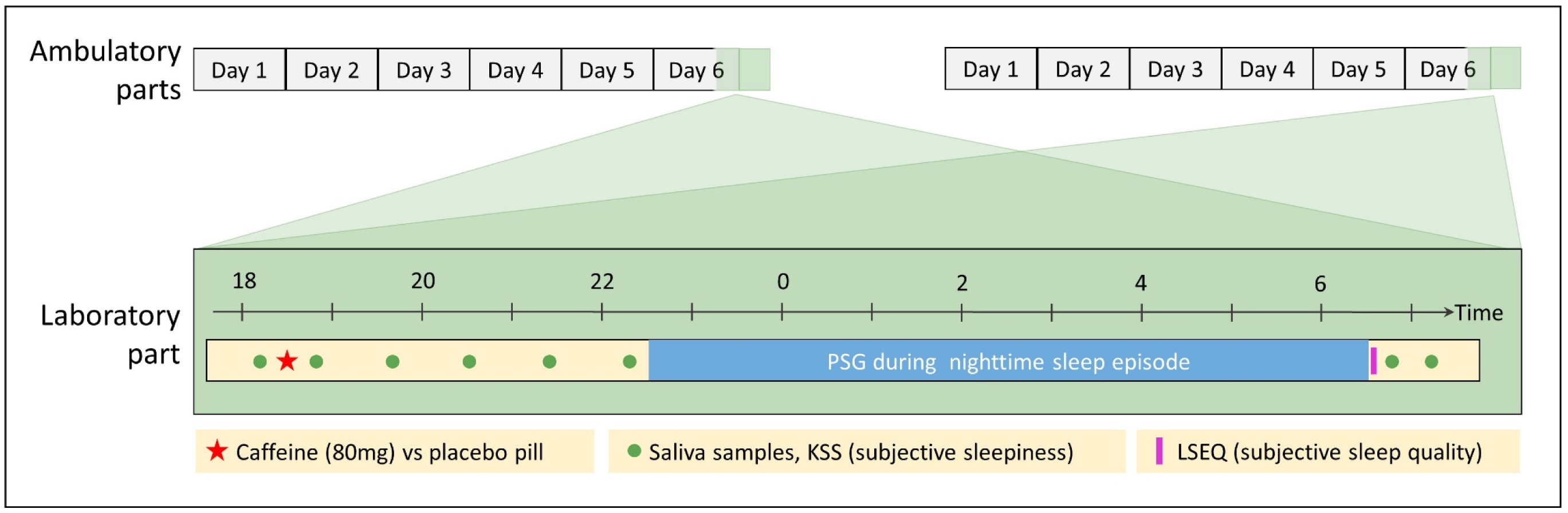
Illustration of study protocol. In each of the two conditions (caffeine and placebo), the protocol started with an ambulatory part of 6 days during which participants were asked to abstain from caffeine and to adhere to a fixed sleep-wake cycle (8h sleep per day, no naps allowed). Sleep-times were adjusted to the participant’s individual and social demands and participants were allowed to deviate by ±1 h from the times set. In the evening of day six of the ambulatory period, the laboratory part started 5 h prior to bedtime (i.e., on average at 17:36). Before and after treatment (caffeine vs. placebo), administered 4 h before habitual bedtime, we assessed subjective sleepiness (by the Karolinska Sleepiness Scale, KSS, (1)) and collected saliva to determine caffeine, paraxanthine and melatonin levels. Polysomnography (PSG) was conducted during the scheduled sleep episode. Subjective sleep quality was measured by the Leeds Sleep Evaluation Questionnaire (LSEQ, (2)) right after awakening.

**Supplemental Table 1.** Amount of sleep, sleep stages and latencies to sleep stages per condition.

| Parameter | Condition |  | Significance |
| --- | --- | --- | --- |
| | Caffeine ($M \pm SD$) | Placebo ($M \pm SD$) | $P$ (two-sided) |
| Total sleep time (min) | 454.56 ± 30.97 | 467.06 ± 8.68 | 0.07 |
| Wake after sleep onset (min) | 16.55 ± 25.71 | 6.60 ± 6.46 | 0.07 |
| Stage 1 (% of TST) | 11.46 ± 4.93 | 8.98 ± 6.48 | 0.02 |
| Stage 2 (% of TST) | 31.98 ± 6.69 | 31.06 ± 9.13 | 0.54 |
| Stage 3 (% of TST) | 36.91 ± 8.69 | 40.51 ± 11.41 | 0.05 |
| Stage REM (% of TST) | 19.66 ± 3.71 | 19.46 ± 3.81 | 0.86 |
| Latency to stage 1 (min) | 8.88 ± 7.38 | 5.57 ± 3.54 | 0.08 |
| Latency to stage 2 (min) | 12.39 ± 8.51 | 8.86 ± 6.05 | 0.15 |
| Latency to stage 3 (min) | 22.33 ± 7.91 | 16.78 ± 8.02 | 0.03 |
| Latency to REM (min) | 118.13 ± 33.36 | 132.63 ± 39.41 | 0.11 |
| Latency to persistent sleep (min) | 10.56 ± 7.54 | 6.70 ± 6.09 | 0.09 |
| Sleep efficiency (%) | 94.70 ± 6.40 | 97.46 ± 1.87 | 0.05 |

Indicators of significance ( $P$ -Values) are derived from t-tests for dependent samples, conducted for each variable separately. Total sleep time (TST): Sum of time in any sleep stage; REM: Rapid eye movement sleep; latency to sleep stages: Time from lights out to the first occurrence of each stage; latency to persistent sleep: Time from lights out to the first occurrence of 10 min in any sleep stage; Sleep efficiency: Total sleep time/Time in bed \* 100.

